# Plasmid diversity among genetically related *Klebsiella pneumoniae bla*_KPC-2_ and *bla*_KPC-3_ isolates collected in the Dutch national surveillance

**DOI:** 10.1101/2020.01.23.917781

**Authors:** Antoni P.A. Hendrickx, Fabian Landman, Angela de Haan, Dyogo Borst, Sandra Witteveen, Marga van Santen, Han van der Heide, Leo M. Schouls, the Dutch CPE surveillance Study Group

## Abstract

Carbapenemase-producing *Klebsiella pneumoniae* emerged over the past decades as an important pathogen causing morbidity and mortality in hospitalized patients. For infection prevention and control, it is important to track the spread of bacterial strains in humans including the plasmids they contain. However, little is known concerning the plasmid repertoire among *K. pneumoniae* strains. Therefore, the major aim was to recapitulate the size, contents and diversity of the plasmids of genetically related *K. pneumoniae* strains harboring the beta-lactamase gene *bla*_KPC-2_ or *bla*_KPC-3_ to determine their dissemination in the Netherlands and the former Dutch Caribbean islands from 2014-2019. Next-generation sequencing was combined with long-read third-generation sequencing to reconstruct 18 plasmids of *K. pneumoniae*. wgMLST revealed five genetic clusters (termed KpnClusters) comprised of *K. pneumoniae bla*_KPC-2_ isolates and four clusters consisted of *bla*_KPC-3_ isolates. Each cluster was characterized by a distinct resistome and plasmidome. KpnCluster-019 *bla*_KPC-2_ isolates were found both in the Netherlands and the Caribbean islands. *K. pneumoniae bla*_KPC-3_ isolates were found in the collection of the Netherlands. The 18 plasmids were mostly unrelated and varied between *K. pneumoniae bla*_KPC-2_ and *bla*_KPC-3_ clusters. However, the large and medium sized plasmids contained a variety of antibiotic resistance genes, transposons, insertion sequence elements, conjugal transfer systems, cation transport systems, toxin/antitoxin systems, and prophage-related sequence elements. The small plasmids carried genes implicated in virulence. Thus, implementing long-read plasmid sequencing analysis for *K. pneumoniae* surveillance provided important insights in the success and understanding of transmission of a KpnCluster-019 *bla*_KPC-2_ strain between the Netherlands and the Caribbean.

**Importance:** Carbapenemase-producing *Klebsiella pneumoniae* has spread globally and is of great concern for debilitated patients. *K*. *pneumoniae* is notorious for spreading antimicrobial resistance genes by plasmids among *Enterobacterales*. Combining short and long read sequencing enables reconstruction of plasmids containing antibiotic resistance genes, conjugation machinery, transposons, toxins and/or virulence determinants and thereby enhancing international pathogen surveillance.

## Introduction

Antimicrobial resistance is spreading rapidly among *Enterobacterales*, including *Klebsiella pneumoniae*, *Escherichia coli* and *Enterobacter* spp. (1). Within the cell, extra-chromosomal DNA such as plasmids encode genes that confer resistance to last resort antibiotics, including carbapenems and colistin, and can transfer between *Enterobacterales* (2). Currently, carbapenemase-producing *Enterobacterales* (CPE) rank among the most problematic nosocomial pathogens with limited outlook on novel effective therapeutics (3, 4). With the current increase of multidrug-resistant infections with CPE worldwide, total healthcare costs are anticipated to increase. *K. pneumoniae* is often referred to as the “canary in the coalmine”, as new antimicrobial resistance (AMR) genes have been associated with *K. pneumoniae* in the first clinical reports prior dispersal of the AMR genes among other Gram-negative bacteria (5). Most newly acquired AMR genes of *K. pneumoniae* are the result of horizontal gene transfer through conjugative plasmids (6–8). The *K. pneumoniae* carbapenemase KPC encoded by the *bla*_KPC_ gene is an Ambler class A serine carbapenemase, which is often located on a transmissible plasmid-associated transposon Tn*4401*, or variants hereof (9–12). Tn*4401* consists of flanking imperfect repeat sequences, a Tn*3* transposase gene, a Tn*3* resolvase gene and the IS*Kpn6* and IS*Kpn7* insertion sequences (10). The *bla*_KPC-2_ and *bla*_KPC-3_ carbapenemases are the most commonly identified variants that have spread globally and provide resistance to penicillins, carbapenems, cephalosporins, cephamycins and monobactams (13, 14). The KPC-2 and KPC-3 carbapenemases differ in only one amino acid as a histidine at position 272 is mutated to tyrosine (H272Y) in the KPC-3 variant (15).

CPE isolates including *K. pneumoniae* are routinely send to the National Institute for Public Health and the Environment (RIVM) and are typed by Illumina next-generation sequencing (NGS) in the Dutch National CPE Surveillance program to identify AMR genes and to determine possible transmission of strains (16). NGS typically yields short sequence reads of 150 bases, thereby hampering the assembly of complete chromosomes and plasmids (17). This is often due to large mobile genetic elements, such as insertion sequence elements, transposons, and other repetitive sequences *e.g.* tandem repeat regions of >1500 bp in size. However, combining Illumina NGS sequencing with long-read third generation sequencing (TGS), which produces 1,000 to 500,000 bases or longer sequence reads, can overcome this problem and enables the reconstruction of chromosomes and complete plasmids (18, 19). Currently, the transmission of *K. pneumoniae* between persons in different countries and the impact hereof is not thoroughly understood. It is also not clear whether plasmids of *K. pneumoniae* circulate endemically in the Netherlands or that are introduced from other high-prevalence countries. While the prevalence of carbapenemase producing *K. pneumoniae* and associated infections in the Netherlands is relatively low, the establishment of genomic surveillance of *K. pneumoniae* using TGS is of high importance (20, 21). It provides for insights in the transmission of specific strains containing plasmids with AMR genes and/or virulence determinants. We therefore investigated the distribution of *K. pneumoniae* cluster isolates harboring *bla*_KPC-2_ or *bla*_KPC-3_ alleles obtained from the Dutch National CPE Surveillance Program and analyzed the contents of its plasmids using long read third-generation sequencing.

## Results

### Distribution and genetic relationship of *bla*_KPC-2_ and *bla*_KPC-3_ carrying *K. pneumoniae*

A collection of 480 carbapenemase-producing *K. pneumoniae* isolates submitted to the Dutch National CPE Surveillance program from January 1^st^ 2014 until June 30^th^ 2019 to the National Institute for Public Health and the Environment (RIVM) were included in this study. The study collection comprised 84 *K. pneumoniae bla*_KPC_-positive isolates of which 51 contained the *bla*_KPC-2_ allele and 33 harbored the *bla*_KPC-3_ allele (Table 1). Sixty isolates originated from the Netherlands and 24 isolates originated from the Caribbean. Of the 24 Caribbean isolates, 22 carried the *bla*_KPC-2_ allele and only two contained the *bla*_KPC-3_ allele. Whole genome multi-locus sequence typing (wgMLST), using an in-house wgMLST scheme based on 4,978 genes, of the 480 carbapenemase-producing *K. pneumoniae* isolates collected in the RIVM revealed that 23 *K. pneumoniae bla*_KPC-2_ isolates grouped together in five distinct genetic clusters. Fifteen *K. pneumoniae bla*_KPC-3_ isolates grouped in four distinct clusters which were obtained from the

**Table 1.** Distribution of *K. pneumoniae bla*_KPC-2_ and *bla*_KPC-3_ isolates and resistance to meropenem. Based on the clinical breakpoints according to EUCAST, the isolates were classified as sensitive (S; <2 mgl/L), intermediate (I; ≥2 to 8 mg/L) and resistant (R; >8 mg/L).

| *bla*<sub>KPC</sub> allele | KpnCluster | wgMLST<br>Cluster type | The Netherlands |  |  | Caribbean |  |  | Total |
| --- | --- | --- | --- | --- | --- | --- | --- | --- | --- |
|  |  |  | S | I | R | S | I | R |  |
| *bla*<sub>KPC-2</sub> | KpnCluster-003 | 2521 |  |  | 2 |  |  |  | 2 |
|  | KpnCluster-005 | 123 |  |  | 3 |  |  |  | 3 |
|  | KpnCluster-019 | 2494 | 1 | 2 |  | 6 | 4 |  | 13 |
|  | KpnCluster-021 | 2588 |  |  |  |  | 1 | 2 | 3 |
|  | KpnCluster-041 | 2494 |  |  |  | 2 |  |  | 2 |
|  | Non-KpnCluster | variant | 3 | 4 | 14 | 1 | 2 | 4 | 28 |
|  | Subtotal |  |  |  |  |  |  |  | 51 |
| *bla*<sub>KPC-3</sub> | KpnCluster-008 | 53 |  |  | 4 |  |  |  | 4 |
|  | KpnCluster-025 | 1257 |  | 1 | 6 |  |  |  | 7 |
|  | KpnCluster-038 | 1752 |  |  | 2 |  |  |  | 2 |
|  | KpnCluster-050 | 1969 | 1 |  | 1 |  |  |  | 2 |
|  | Non-KpnCluster | variant | 3 | 5 | 8 | 1 |  | 1 | 18 |
|  | Subtotal |  |  |  |  |  |  |  | 33 |
| Total |  |  | 8 | 12 | 40 | 10 | 7 | 7 | 84 |

Netherlands and 46 isolates were unrelated. The *K. pneumoniae* cluster isolates (termed KpnClusters) had unique wgMLST cluster types, and were not described previously (Table 1, Fig. 1). KpnCluster-003 and KpnCluster-005 were comprised of five *K. pneumoniae bla*_KPC-2_ isolates that were exclusively obtained from the Netherlands, while KpnCluster-021 and KpnCluster-041 contained five isolates from the Caribbean. The majority (*n* = 10) of the KpnCluster-019 isolates were obtained from the Caribbean. However, three isolates were from the collection of the Netherlands. One person from whom a KpnCluster-019 isolate was retrieved in August 2017 in the Netherlands, lived in the Caribbean until June 2017 and migrated to the Netherlands in July, demonstrating intercontinental transmission. No epidemiological data could be retrieved from the other two Dutch KpnCluster-019 isolates. Furthermore, most genetic clusters were only distantly related to each other (Fig. 1). The genetic distance between KpnCluster-019 and KpnCluster-041 was 30 alleles and for KpnCluster-003 and KpnCluster-005 53 alleles. KpnCluster-008 differed 132 alleles from KpnCluster-005. While the allelic difference between these clusters was low, the other genetic clusters differed 3573 to 3610 alleles from KpnCluster-005. This confirmed that most clusters were unrelated, and it is in line with the location of these genetic clusters in the minimum spanning tree.

**Fig. 1.**
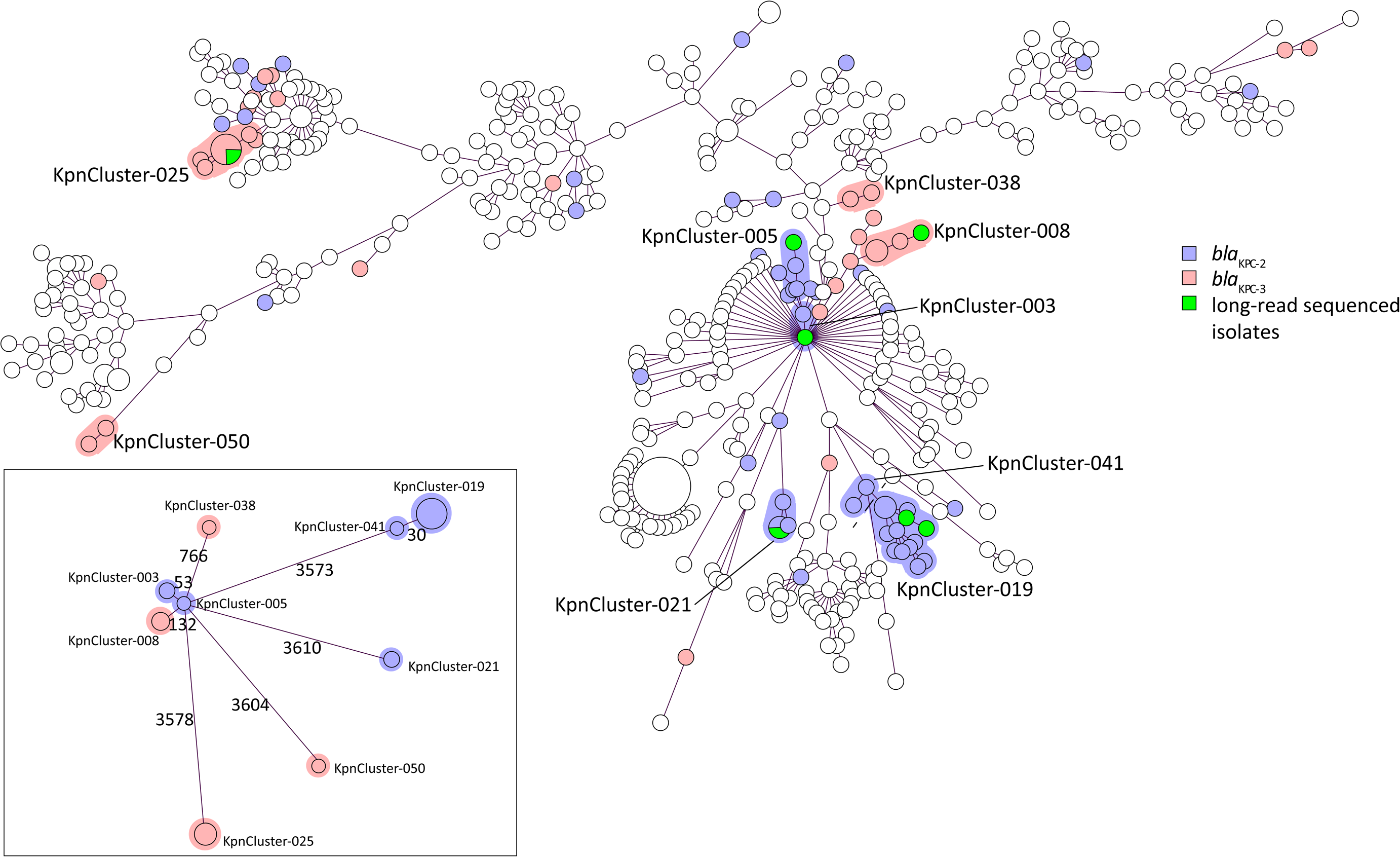
Minimum spanning tree based on wgMLST of 480 sequenced *K. pneumoniae* isolates. Circles represent *K. pneumoniae* isolates, and the sizes of the circles indicate the numbers of isolates. Lines connecting the circles represent the genetic distance in numbers of alleles; the longer the connecting line, the larger the genetic distance. *K. pneumoniae bla*_KPC-2_ isolates were marked blue and *K. pneumoniae bla*_KPC-3_ were marked magenta. *K. pneumoniae bla*_KPC-2_ or *bla*_KPC-3_ cluster isolates that were sequenced with TGS were marked green. Genetic clusters were indicated with either a blue or a magenta halo around the circles, if two or more isolates differ ≤ 20 alleles. A categorical coefficient was used for the clustering. Cluster names are indicated. Inset: genetic distance between the KpnClusters in which the allelic difference is indicated by numbers.

### The resistome diversity among genetic clusters

Analysis of the NGS-derived resistomes of the cluster and non-cluster isolates showed that *K. pneumoniae* harbored either the *bla*_KPC-2_ or the *bla*_KPC-3_ allele, none of the isolates carried both alleles (Fig. 2, Suppl. Fig. 1). Nearly all of the *K. pneumoniae* isolates contained the *fosA*, *oqxA* and *oqxB* genes conferring resistance to fosfomycin and fluoroquinolone antibiotics, respectively. An unweighted hierarchical clustering (UPGMA) based on the presence or absence of AMR genes revealed that most genetic cluster isolates group together per cluster, since the resistomes were more than 85% similar. In contrast to this, the resistomes of the non-cluster isolates were very diverse and less related since the resistomes of these isolates were less than 85% similar (Suppl. Fig. 1). Likewise, the resistomes of one group of *K. pneumoniae* KpnCluster-003 *bla*_KPC-2_ and KpnCluster-008 *bla*_KPC-3_ cluster isolates with 53 to 132 alleles difference were also unrelated. KpnCluster-019 isolates are unique when compared to the *bla*_KPC-2_ clusters KpnCluster-003, KpnCluster-005, and KpnCluster-021, in that they carried aminoglycoside (*aac*(*3*)*-IIa*), extended spectrum beta-lactams (*bla*CTX-M-15, *bla*SHV-26), fluoroquinolone (*qnrB1*) and tetracyclin (*tetA*) antimicrobial resistance (AMR) genes. KpnCluster-019 and KpnCluster-041 isolates, obtained from the Caribbean, were closely related based on wgMLST, and group together based on the resistome too. The absence of AMR genes *aph(3’’)-Ib*, *aph*(*6*)*-Id* and *sul2* in five of KpnCluster-019 isolates, including the TGS sequenced isolates, indicate the absence of an AMR gene containing plasmid. In addition, the presence of three KpnCluster-019 isolates from the Netherlands with varying resistomes within the cluster suggests additional transmissions. KpnCluster-025 *bla*_KPC-3_ isolates contained the aminoglycoside (*aac*(*3*)*-IIa*) and beta-lactam AMR genes (*bla*SHV-28), while the other Kpn *bla*_KPC-3_ clusters did not. Notably, *mcr* genes conferring resistance to colistin were not detected in the 84 isolates analyzed. The majority of the *K. pneumoniae bla*_KPC-2_ and *bla*_KPC-3_ isolates were resistant to meropenem (47/84; 56%). More specifically, seven of the 23 *K. pneumoniae bla*_KPC-2_ cluster isolates (30%) and 13 of the 15 *bla*_KPC-3_ cluster isolates (87%) were resistant to meropenem. The remainders of the cluster and non-cluster isolates were intermediate resistant or sensitive for meropenem (Table 1).

**Fig. 2.**
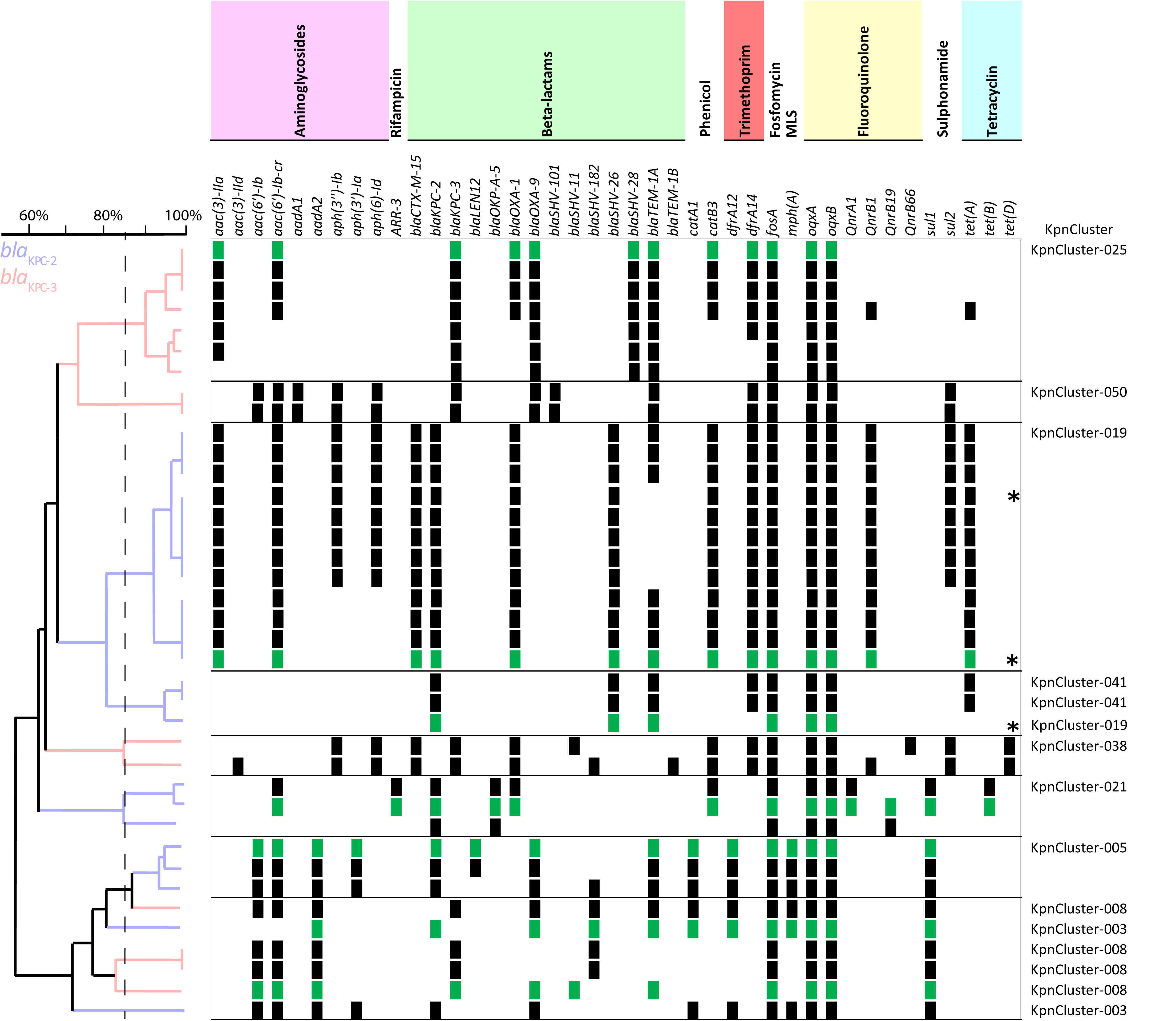
Resistome of *K. pneumoniae bla*_KPC-2_ and *bla*_KPC-3_ cluster isolates. *K. pneumoniae bla*_KPC-2_ and *bla*_KPC-3_ cluster isolates were indicated on the y-axis and AMR genes on the x-axis. Antibiotic classes are indicated above the AMR genes in different colors. The clustering was based on the presence (squares) and absence of AMR genes. Resistance genes in *K. pneumoniae bla*_KPC-2_ or *bla*_KPC-3_ cluster isolates that were sequenced with TGS were marked with green squares. Genetic relatedness was depicted in an UPGMA tree in which *K. pneumoniae bla*_KPC-2_ isolates were marked with blue branches, and *K. pneumoniae bla*_KPC-3_ were marked magenta. Dutch KpnCluster-019 isolates were marked with an *. A dotted line marks the 85% cut off.

### Antibiotic resistance genes among the genomic elements of the distinct genetic clusters

Long-read sequencing of seven isolates from six of the nine genetic *K. pneumoniae bla*_KPC_ clusters, revealed 18 plasmids with varying sizes (Fig. 3). Plasmids containing either the *bla*_KPC-2_ or *bla*_KPC-3_ allele were diverse in size. The large (≥150-250 kb) and medium (≥50-150 kb) sized plasmids contained one or two replicons from the incompatibility group IncFIB(K) and IncFII(K), IncHI2 and IncHI2a, or IncFIB(pQil) (Fig. 3). The small plasmids (<50 kb) contained ColRNAI or IncX3/IncL/IncP6 type of replicons. The chromosomes of the analyzed isolates contained on average five acquired AMR genes, while the plasmids contained on average nine AMR genes. Fourteen of the 18 plasmids contained AMR genes from various classes and four plasmids from the isolate of KpnCluster-021 did not. The AMR genes conferring resistance to phenicol, trimethoprim and macrolide antibiotics were located only on medium or large sized plasmids. The small plasmids had one or two AMR genes conferring resistance to aminoglycosides or beta-lactams. Resistance genes for fosfomycin (*fosA*) and fluoroquinolones (*oqxA* and *oqxB*) were exclusively located on the chromosomes of the seven cluster isolates. KpnCluster-019 and KpnCluster-021 associated with the Caribbean contained plasmids encoding genes for phenicol and tetracyclin resistance. The KpnCluster-019 and KpnCluster-021 plasmids were not found in non-cluster isolates, whereas the plasmids of the other clusters were detected in a subset non-cluster isolates (Fig. 3). The plasmids of KpnCluster-003 and KpnCluster-005 were present in each of its cluster isolates, however, in isolates of the other clusters occasionally plasmids were lost, thereby impacting the composition of the resistome (Fig. 2 and 3).

**Fig. 3.**
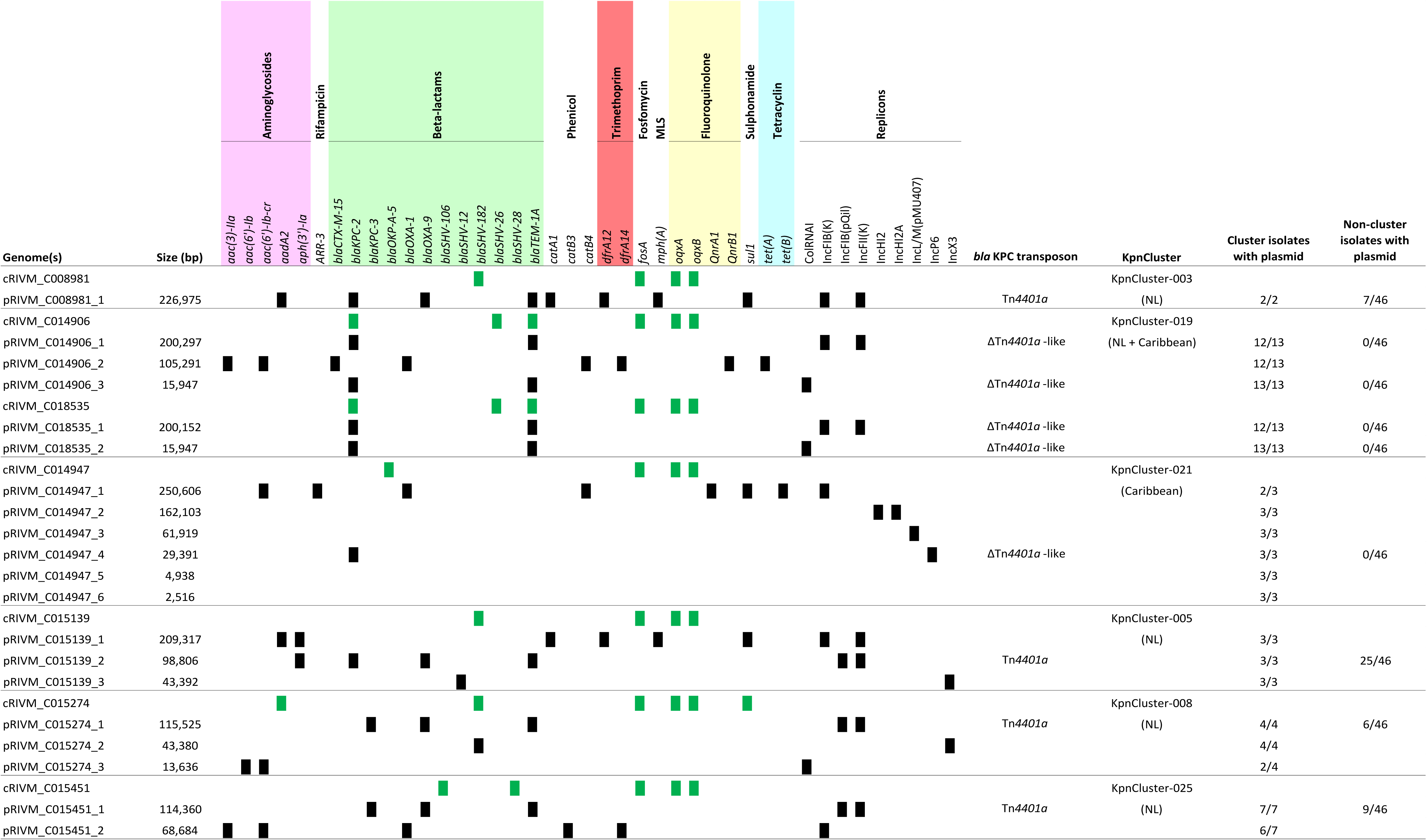
Antimicrobial resistance genes on chromosomes and plasmids. The presence (black squares) and absence of AMR genes among the chromosomes and 18 plasmids of seven TGS sequenced isolates (green). Chromosomes (cRIVM_C0xxxx) and plasmids (pRIVM_C0xxxx) are depicted on the Y-axis, and AMR genes on the x-axis. Antibiotic classes are indicated above the AMR genes in different colors.

The *bla*_KPC-2_ KpnCluster-019 isolates were obtained from both the Caribbean and the Netherlands, while *bla*_KPC-2_ KpnCluster-021 isolates originated only from the Caribbean (Table 1, Fig. 3). In the KpnCluster-019 isolate RIVM_C014906, three copies of the bla_KPC-2_ gene were present, while other cluster isolates had only one *bla*_KPC_ copy. One copy was located in the chromosome, one copy in the 200 kb plasmid pRIVM_C014906_1 and a third copy on the 16 kb plasmid pRIVM_C014906_3. All these three *bla*_KPC-2_ copies were located on a highly similar *Tn4401*a-derived Δ*Tn4401*a-like transposon of 5.6 kb in this strain. The chromosomes contained this Δ*Tn4401*a-like transposon in the exact same region. KpnCluster-003, KpnCluster-005, KpnCluster-008 and KpnCluster-025 consist of isolates that were obtained in the Netherlands and in these isolates the *bla*_KPC_ allele was located on a Tn*4401a* transposon of 10 kb.

### Comparison of the *K. pneumoniae* plasmid content

An UPGMA clustering based on the DNA sequence of the 18 plasmids revealed that the majority of the plasmids were unrelated (Fig. 4). The largest two plasmids pRIVM_C008981_1 and pRIVM_C014947_1 carried the largest number of genes and this number decreased by the decreasing size of the plasmids. Most the plasmid located genes had unknown function. The large and medium sized plasmids contained the *klcA* gene, encoding an antirestriction protein implicated in the facilitation of *bla*_KPC_ allele transfer (22). None of the plasmids contained known virulence determinants such as *rmpA*, *rmpA2*, *iroBC*, or *iucABC* implicated in hypervirulence (23, 24). Comparison of the large plasmids revealed that pRIVM_C008981_1 and pRIVM_C015139_1 from KpnCluster-003 and KpnCluster-005 displayed 90% similarity (Fig. 4). Plasmid pRIVM_C014947_1 was not related to any other of the large plasmids. Despite the low similarity, these large plasmids shared important clusters of genes among them. They all contained the *silE* and *silP* genes encoding a silver-binding protein and a silver exporting ATPase, *cusSRCFB* genes implicated in cation efflux, the *copABCD-pcoE* genes involved in copper resistance and the *arsHACBAD* arsenic resistance gene cluster. These large plasmids also contained *fecIRABCDE* implicated in Fe(3+)-dicitrate transport, the *traIDSQCVAJM-ylpA* plasmid conjugation gene cluster, and the *higA-higA1* antitoxins, except pRIVM_C014947_1 and pRIVM_C014947_2. In addition, the large plasmids also contained a proportion of plasmid-specific and thus *K. pneumoniae* cluster specific content (Suppl. Fig. 2).

**Fig. 4.**
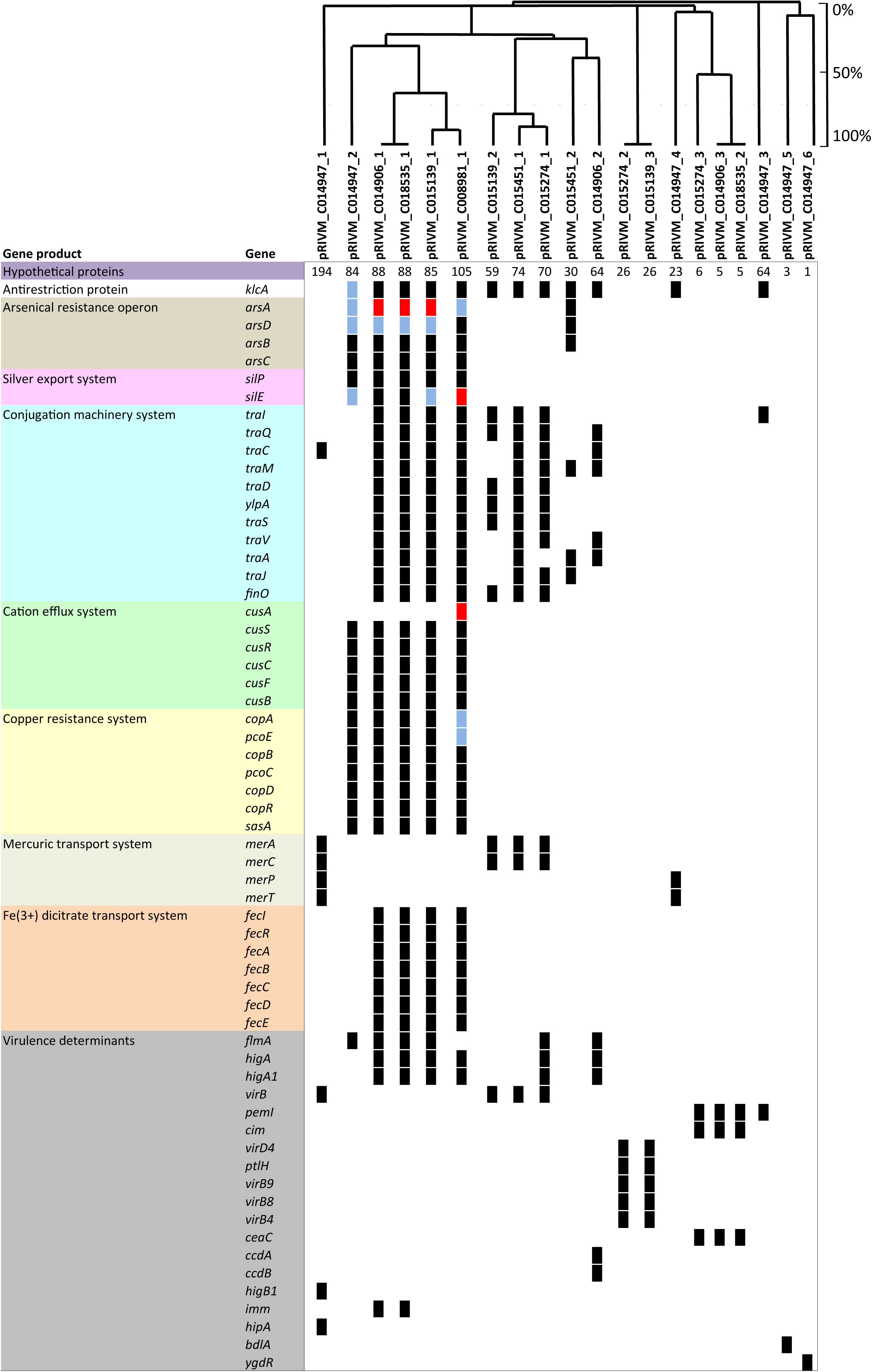
*K. pneumoniae* plasmid gene content. An UPGMA clustering was performed based on the plasmid DNA sequence for the determination of the genetic relation among the 18 plasmids. Similarity is indicated on the y-axis using a scale from 0 (not similar) to 100% (identical). A similarity of ≥85 to 100% is regarded as the same plasmid. The plasmids are indicated on the x-axis. The presence (black squares) and absence is indicated of annotated genes among the 18 plasmids of seven TGS sequenced isolates. If a gene was present twice, blue squares were used and more than 2, red squared were used. Colors indicated different groups of genes with a specific function.

The medium-sized plasmids contained the *virB* virulence regulon transcriptional activator and the *merAC* mercuric reductase and transport protein. While pRIVM_C015274_1 and pRIVM_C015451_1 contained a plasmid conjugation gene cluster, pRIVM_C014906_2 and pRIVM_C015139_2 contained truncated versions hereof. The more distantly related pRIVM_C014906_2 plasmid from KpnCluster-019 had in addition to the *higA-higA1* antitoxins also a *ccdA-ccdB* toxin-antitoxin system. The small plasmids (<50 kb) contained genes implicated in virulence. Plasmids pRIVM_C015139_3 and pRIVM_C015274_2 displayed 99% similarity and carried the *virD4-B9-B8-B4*-*ptlH* Type IV secretion system. pRIVM_C014947_4 contained a *merPT* mercuric transport system, while pRIVM_C014906_3 and pRIVM_C015274_3 carried a *ceaC* colicin-E3. The plasmid pRIVM_C014947_5 contained the *bdlA* gene encoding a biofilm dispersion protein.

### Transposable elements in *K. pneumoniae* plasmids

The large and medium sized plasmids contained the most transposase sequences, and each plasmid had its unique transposon signature (Fig. 5). The IS1 and IS3 transposase families dominated in the *K. pneumonia*e plasmids. The IS1 family transposase was found most frequently among the plasmids and in most copies within plasmids. In the large and medium sized plasmids, the *bla*_KPC_ allele was located on a Tn*4401a* transposon, except in pRIVM_C014906_1. In the small plasmids carrying a *bla*_KPC_, the carbapenemase allele was located on a Δ*Tn4401*a-like transposon. The large plasmids pRIVM_C008981_1, pRIVM_C015139_1 and pRIVM_C014906_1 harbored 37, 32 and 31 annotated tranposases, respectively. In contrast, the largest plasmid pRIVM_C014947_1 of 250.6 kb from KpnCluster-021 contained only 12 transposons. The remainder of the plasmids from KpnCluster-021 also contained very few transposase sequences, in contrast to the other plasmids from the different clusters. The highly related pRIVM_C015139_3 and pRIVM_C015274_2 plasmids (99% similarity) had identical transposons. While IS66 and IS110 family transposase sequences also dominate in the large plasmids, the medium sized plasmids contained IS3 family type of transposases. The medium sized plasmids contained eleven to 23 transposases, and the small plasmids less than ten.

**Fig. 5.**
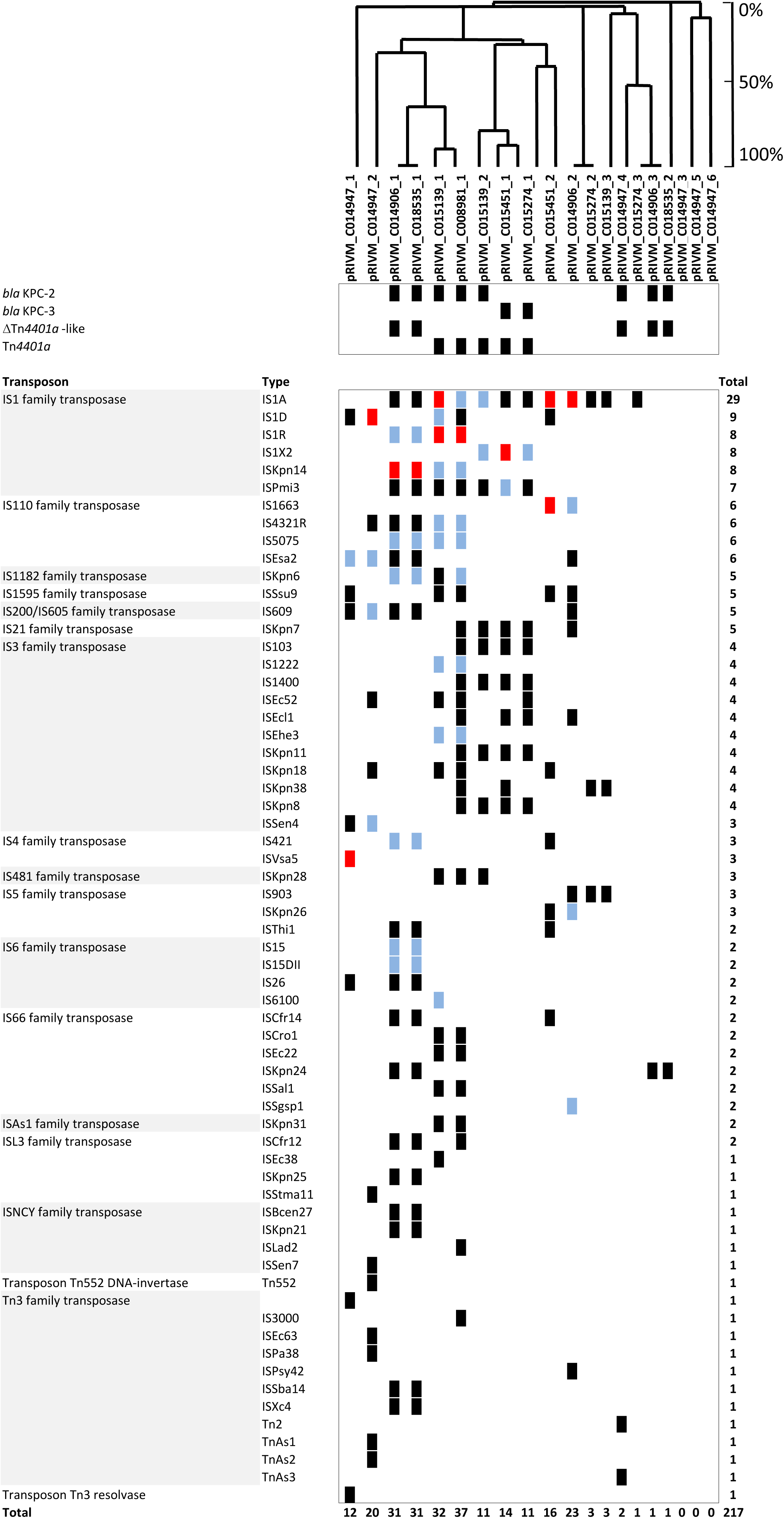
*K. pneumoniae* plasmid-localized transposases. The presence (black squares) and absence is indicated of annotated transposases among the 18 plasmids of six TGS sequenced isolates. The plasmids are indicated on the x-axis. If a transposon was present twice, blue squares were used and more than 2, red squared were used. The light grey area indicates specific transposons found in only one plasmid.

### Similarity with previously reported plasmids

BLAST analysis of the *K. pneumoniae* plasmids identified in this study showed that 8 of the 18 plasmids were similar to previously reported plasmids in the NCBI sequence database (Table 2). These plasmids covered five distinct genetic clusters, except pRIVM_C008981_1 from KpnCluster-003. To date, none of these plasmids were reported to be implicated in healthcare-associated outbreaks. Plasmids pRIVM_C008981_1, pRIVM_C014906_1, pRIVM_C014906_3 containing *bla*_KPC-2_ and pRIVM_C015274_1 harboring *bla*_KPC-3_ from distinct genetic clusters only had low sequence coverage 35-87% with plasmids present in the NCBI sequence database. The other *bla*_KPC-2_ and *bla*_KPC-3_ plasmids had high (93-99%) sequence coverage, indicating that these similar plasmids were detected previously by other researchers. Plasmids pRIVM_C014906_2, pRIVM_C015139_1, pRIVM_C015274_2 and pRIVM_C015274_3, not carrying a *bla*_KPC_ allele, displayed 97-100% sequence coverage and 99-100% identity to plasmids isolated from *K. pneumoniae* from different countries (Table 2). Plasmids pRIVM_C014947_5 and pRIVM_C014947_6 from KpnCluster-021 had 100% sequence coverage with 92.18 to 99.99% identity with plasmids isolated from *Enterobacter hormaechei*. Plasmids similar to pRIVM_C014947_1, pRIVM_C014947_3, pRIVM_C014947_5 and pRIVM_C014947_6 from KpnCluster-021 were detected previously in a variety of hosts, *e.g. Salmonella enterica*, *K. pneumoniae*, and *E. hormaechei*, suggesting these plasmids are broad-host range.

**Table 2.**
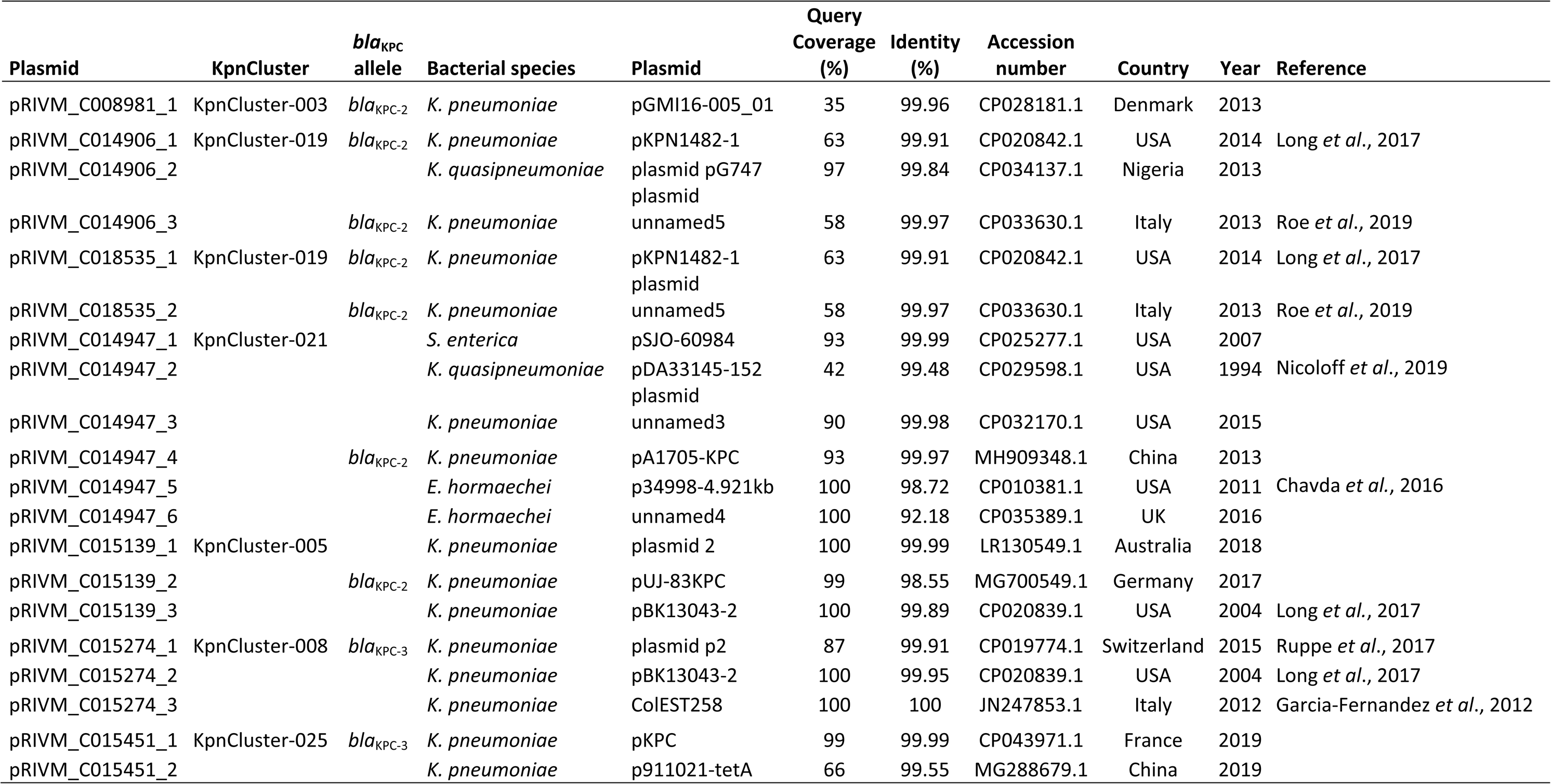
BLAST similarity analysis of *K. pneumoniae* plasmids.

| Plasmid | KpnCluster | *bla*<sub>KPC</sub><br>allele | Bacterial species | Plasmid | Query<br>Coverage<br>(%) | Identity<br>(%) | Accession<br>number | Country | Year | Reference |
| --- | --- | --- | --- | --- | --- | --- | --- | --- | --- | --- |
| pRIVM_C008981_1 | KpnCluster-003 | *bla*<sub>KPC-2</sub> | *K. pneumoniae* | pGMI16-005_01 | 35 | 99.96 | CP028181.1 | Denmark | 2013 |  |
| pRIVM_C014906_1 | KpnCluster-019 | *bla*<sub>KPC-2</sub> | *K. pneumoniae* | pKPN1482-1 | 63 | 99.91 | CP020842.1 | USA | 2014 | Long *et al.*, 2017 |
| pRIVM_C014906_2 |  |  | *K. quasipneumoniae* | plasmid pG747 | 97 | 99.84 | CP034137.1 | Nigeria | 2013 |  |
| pRIVM_C014906_3 |  | *bla*<sub>KPC-2</sub> | *K. pneumoniae* | unnamed5 | 58 | 99.97 | CP033630.1 | Italy | 2013 | Roe *et al.*, 2019 |
| pRIVM_C018535_1 | KpnCluster-019 | *bla*<sub>KPC-2</sub> | *K. pneumoniae* | pKPN1482-1<br>plasmid | 63 | 99.91 | CP020842.1 | USA | 2014 | Long *et al.*, 2017 |
| pRIVM_C018535_2 |  | *bla*<sub>KPC-2</sub> | *K. pneumoniae* | unnamed5 | 58 | 99.97 | CP033630.1 | Italy | 2013 | Roe *et al.*, 2019 |
| pRIVM_C014947_1 | KpnCluster-021 |  | *S. enterica* | pSJO-60984 | 93 | 99.99 | CP025277.1 | USA | 2007 |  |
| pRIVM_C014947_2 |  |  | *K. quasipneumoniae* | pDA33145-152<br>plasmid | 42 | 99.48 | CP029598.1 | USA | 1994 | Nicoloff *et al.*, 2019 |
| pRIVM_C014947_3 |  |  | *K. pneumoniae* | unnamed3 | 90 | 99.98 | CP032170.1 | USA | 2015 |  |
| pRIVM_C014947_4 |  | *bla*<sub>KPC-2</sub> | *K. pneumoniae* | pA1705-KPC | 93 | 99.97 | MH909348.1 | China | 2013 |  |
| pRIVM_C014947_5 |  |  | *E. hormaechei* | p34998-4.921kb | 100 | 98.72 | CP010381.1 | USA | 2011 | Chavda *et al.*, 2016 |
| pRIVM_C014947_6 |  |  | *E. hormaechei* | unnamed4 | 100 | 92.18 | CP035389.1 | UK | 2016 |  |
| pRIVM_C015139_1 | KpnCluster-005 |  | *K. pneumoniae* | plasmid 2 | 100 | 99.99 | LR130549.1 | Australia | 2018 |  |
| pRIVM_C015139_2 |  | *bla*<sub>KPC-2</sub> | *K. pneumoniae* | pUJ-83KPC | 99 | 98.55 | MG700549.1 | Germany | 2017 |  |
| pRIVM_C015139_3 |  |  | *K. pneumoniae* | pBK13043-2 | 100 | 99.89 | CP020839.1 | USA | 2004 | Long *et al.*, 2017 |
| pRIVM_C015274_1 | KpnCluster-008 | *bla*<sub>KPC-3</sub> | *K. pneumoniae* | plasmid p2 | 87 | 99.91 | CP019774.1 | Switzerland | 2015 | Ruppe *et al.*, 2017 |
| pRIVM_C015274_2 |  |  | *K. pneumoniae* | pBK13043-2 | 100 | 99.95 | CP020839.1 | USA | 2004 | Long *et al.*, 2017 |
| pRIVM_C015274_3 |  |  | *K. pneumoniae* | ColEST258 | 100 | 100 | JN247853.1 | Italy | 2012 | Garcia-Fernandez *et al.*, 2012 |
| pRIVM_C015451_1 | KpnCluster-025 | *bla*<sub>KPC-3</sub> | *K. pneumoniae* | pKPC | 99 | 99.99 | CP043971.1 | France | 2019 |  |
| pRIVM_C015451_2 |  |  | *K. pneumoniae* | p911021-tetA | 66 | 99.55 | MG288679.1 | China | 2019 |  |

### Prophage sequences in the *K. pneumoniae* cluster genomes

PHASTER analysis revealed that the majority of the large and medium-sized plasmids from different genetic clusters with IncFIB(K) or IncFIB(pQil) and IncFII(K) replicons contained one to four regions with prophage-related sequences *e.g.* genes encoding putative phage integrase, phage-like proteins, coat proteins, and/or tail shaft proteins (Table 3). The size of the prophage sequence regions varied per plasmid. The most commonly found prophage-related sequence in large and medium-sized plasmids of cluster isolates was an *Escherichia* phage RCS47 (Table 3). This sequence entails the 14.2 kb *ygbMLKJI-bla*SHV*-recF-lacY* region flanked by IS26 elements and representing 12% of the RCS47 prophage genome. The small plasmids of <50 kb lacked phage-related sequences. In contrast, the chromosomes of cluster isolates carried at least three to nine phage sequence regions covering 10-50% of the phage genome. These phage sequence regions covered a wide variety of distinct phages, including prophage sequences from *Salmonella*, *Klebsiella*, *Cronobacter*, *Enterobacteria* phages (Suppl. Table 2). The most commonly found prophage sequence in *Klebsiella* chromosomes was the Enterobacteria phage P4.

**Table 3.**
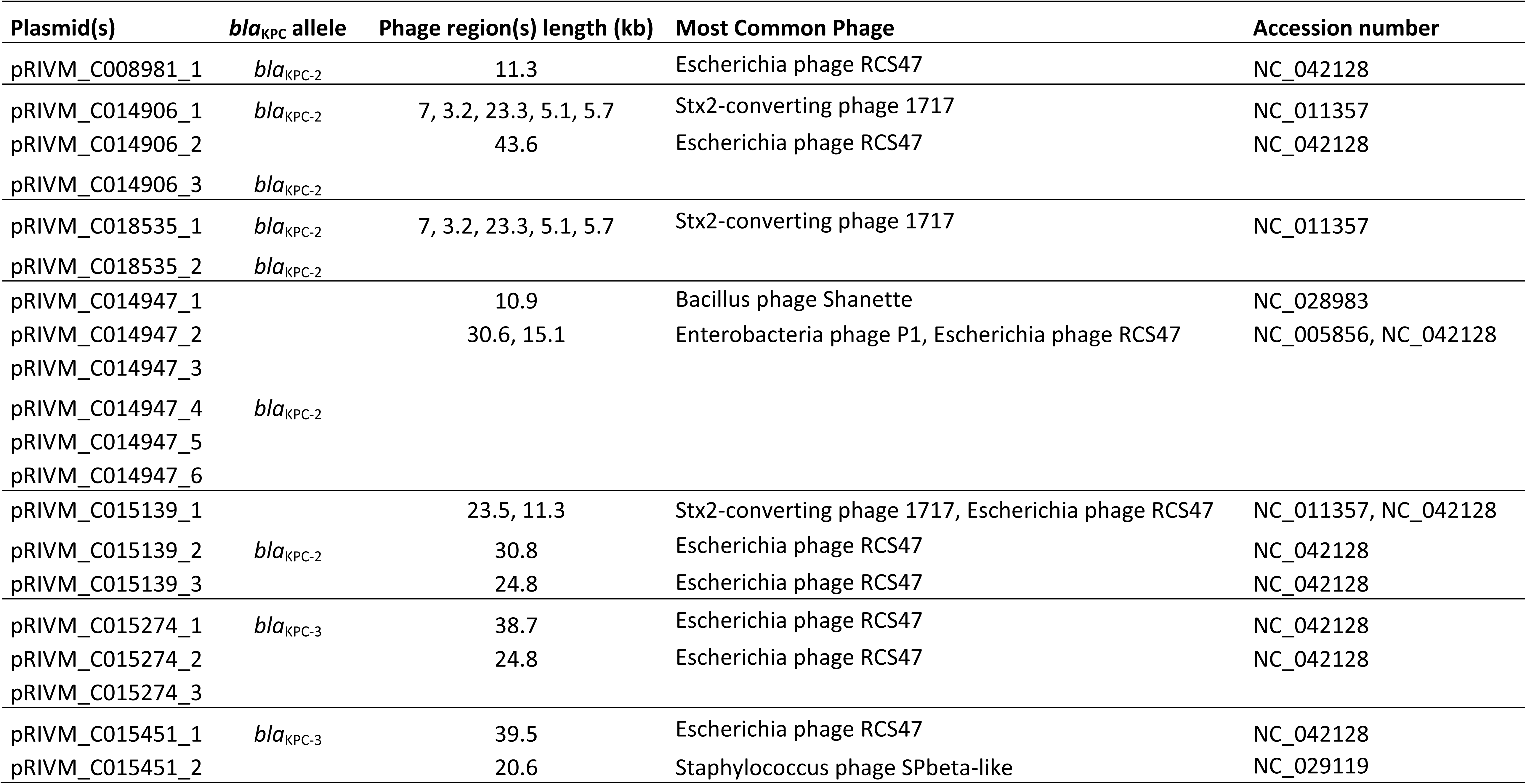
Predicted prophage sequenc 447 es among K. pneumoniae plasmids.

## Discussion

We showed that a *K. pneumoniae* strain carrying *bla*_KPC-2_ was transmitted between the Netherlands and the Caribbean. This is based on the high genetic relatedness of the 13 isolates from KpnCluster-019 as assessed by wgMLST and their highly similar resistome and plasmidome. We found that one person lived in the Caribbean and migrated to the Netherlands. After migration, a KpnCluster-019 isolate was obtained from this person in a Dutch hospital. Possibly other transmissions by other persons could have occurred, but these were not confirmed in this study. By combining short-read with long-read sequencing data, we identified 18 plasmids of seven *K. pneumoniae* isolates from six distinct genetic clusters found in the Netherlands and the Caribbean and analyzed these plasmids for its AMR gene profile, *bla*_KPC_ transposons, replicons, transposon families, and gene content. The plasmid composition varied among the genetic clusters. The cluster isolates had unique wgMLST cluster types which were not published previously and differ from globally circulating extensively drug-resistant (XDR) and highly virulent (hvKp) *K. pneumoniae* strains (23, 24). KpnCluster-019 is unique compared to the other cluster isolates analyzed in this study for the following reasons. First, KpnCluster-019 harbors a unique and extensive set of AMR genes on the chromosome and in its plasmids. Secondly, KpnCluster-019 isolates were the only to contain three copies of the *bla*_KPC-2_ allele, two on two different plasmids and one in the chromosome. The localization of *bla*_KPC-2_ on the chromosome and additional *bla*_KPC-2_ copies have been reported previously and is further complicating the understanding of transmission of multidrug-resistant *K. pneumoniae* (25, 26). Thirdly, KpnCluster-019 and also KpnCluster-021 isolates from the Caribbean harbored the *bla*_KPC-2_ allele on a 5.6 kb Δ*Tn4401*a-like transposon, while the other isolates from the other genetic clusters from the Netherlands contained *bla*_KPC_ on a 10kb Tn*4401a* transposon. Most global descriptions of *K. pneumoniae bla*_KPC_ the past decade have been associated with Tn*4401a* or isoforms hereof (9). The traditional association of *bla*_KPC_ with the Tn*4401a* transposon has possibly been eroded in *K. pneumoniae* isolates from the Caribbean to a smaller variant. This is the first report of identification of a 5.6 kb Δ*Tn4401*a-like *bla*KPC-2 transposon of *K. pneumoniae* in the Netherlands. Preliminary surveillance data analysis revealed that the Δ*Tn4401*a-like element carrying *bla*_KPC-2_ and smaller variants disseminated among *E. cloacae*, *Serratia marcesens*, *K. oxytoca* and *E. coli* in the Netherlands (unpublished data). Future work will seek to understand the dissemination of the Δ*Tn4401*a-like *bla*_KPC-2_ element among CPE in the Netherlands. Lastly, the plasmids of KpnCluster-019 isolates contained also unique plasmid content, including a distinct transposon signature, two toxin-antitoxin systems and a *ceaC* colicin which possibly contribute to the success in survival, niche adaptation or transmission of this strain.

The *K. pneumoniae bla*_KPC-3_ isolates had higher MICs for meropenem than the *K. pneumoniae bla*KPC-2 isolates, which is in line with a previous study (21). The KPC-2 enzyme differs in a single amino acid substitution (Histidine 272 to Tyrosine) from KPC-3. Additional changes in KPC-3 can lead to increased resistance for ceftazidime and cephamycin (27). The increase in meropenem resistance observed in our study is possibly correlated with improved ability of KPC-3 enzymes to hydrolyze the meropenem antibiotic (15). Alternatively, additional beta-lactamase genes such as *bla*_OXA-1_, *bla*_OXA-9_ or *bla*_TEM-1A_ may contribute to increased resistance for meropenem (28).

Despite the limited number of long-read sequenced isolates, we have highlighted important new insights in the genomic surveillance of a notorious multi-antibiotic resistant nosocomial pathogen. In some clusters, the plasmidome varied as this was likely due to loss of a plasmid. Also, the resistome data suggest the presence of other plasmids in cluster isolates that were not present in the isolates that were sequenced using TGS. To overcome this limitation, all isolates used in this study should have been sequenced using long-read third generation sequencing. Nevertheless, we identified plasmids in *K. pneumoniae bla*_KPC-2_ and *bla*_KPC-3_ cluster isolates which vary in size from large, medium and small. The large and medium sized plasmids were enriched for a variety of transposons, conjugation transfer systems, cation efflux systems including Fe(3+)-dicitrate transport, and genes encoding for silver, copper and arsenic resistance. The small plasmids contained putative virulence determinants. The presence of these systems may contribute to the success of transmission of specific *K. pneumoniae* strains in the hospital setting or the community (13, 29, 30). Escherichia RCS47 prophage sequences were found on medium and large plasmids in the cluster isolates analyzed. In contrast, the chromosomes contained a variety of prophage-related sequences. RCS47 is a P1-like bacteriophage carrying the ESBL-encoding *bla*SHV-2 gene was isolated from a clinical *Escherichia coli* strain (31). The prevalence of P1-like prophages in animal and human *E. coli* strain collections was 12.6% (31). The presence of P1-like phage sequences in plasmids of a snapshot of the *K. pneumoniae* population in the Netherlands and the Caribbean suggest that the role of P1-like phages in disseminating antibiotic resistance may be underestimated (32).

In conclusion, long-read sequencing contributed to the understanding of the successful transmission of the KpnCluster-019 *K. pneumoniae bla*_KPC-2_ strain. Plasmid content such as conjugation machinery, transposons, virulence determinants and phages may contribute to diversification, and dissemination of plasmids containing AMR genes, and therefore represent important plasmid features that warrants future investigation. More long-read plasmid sequencing efforts of CPE and *K. pneumoniae* in particular are required to identify the complete plasmid reservoir involved in the spread of antibiotic resistance determinants in the Netherlands and the Caribbean islands.

## Materials and methods

### Bacterial isolates

For the Dutch National carbapenemase-producing Enterobacterales (CPE) Surveillance program, medical microbiology laboratories from the Netherlands andthe Caribbean routinely send CPE isolates with a meropenem minimum inhibitory concentration (MIC) of ≥0.25 µg/ml and/or an imipenem MIC of ≥1 µg/ml or phenotypic (CIM-test) or genotypical evidence of carbapenemase production to the National Institute of Public Health and the Environment (RIVM) (16). For this study, 84 carbapenemase-producing *K. pneumoniae* isolates carrying either the *bla*_KPC-2_ allele or the *bla*_KPC-3_ allele were included and collected in the period from January 1^st^ 2014 until June 30^th^ 2019. Only the first *K. pneumoniae* isolate per patient in this study period was selected. The 84 isolates were obtained from 84 persons and from various isolation sites, *i.e*. rectum/perineum (*n* = 43), throat (*n=* 11), pus (*n* = 2), sputum (*n* = 4), urine (*n* = 10), wound (*n*= 5) and nine were from miscellaneous isolation sites. All bacterial strains were grown aerobically at 37°C on Columbia sheep blood agar plates.

### Antimicrobial susceptibility testing

Resistance to carbapenem was confirmed by assessing the MICs for meropenem for all the 84 isolates using an Etest (bioMérieux Inc., Marcy l’Etoile, France). Based on the clinical breakpoints according to EUCAST, the *K. pneumoniae* isolates were classified as sensitive (≤2 mgl/L), intermediate (>2 mg/L and ≤8 mg/L) and resistant (>8 mg/L) to meropenem. In addition, all isolates were analyzed for the production of carbapenemase using the carbapenem inactivation method (CIM) as described previously (33).

### Next-generation sequencing and wgMLST

All 84 *K. pneumoniae* isolates were subjected to next-generation sequencing (NGS) using the Illumina HiSeq 2500 (BaseClear, Leiden, the Netherlands). The NGS data of the *K. pneumoniae* isolates were used for wgMLST analyses using the in-house wgMLST scheme in SeqSphere software version 6.0.2 (Ridom GmbH, Münster, Germany). Ridom wgMLST cluster nomenclature were depicted in Table 1. The resulting data was imported into Bionumerics version 7.6.3 for subsequent comparative analyses (Applied Maths, Sint-Martens-Latem, Belgium). The antibiotic resistance gene profile and plasmid replicon compositions in all of the isolates were determined by interrogating the online ResFinder (version 3.1.0) and PlasmidFinder (version 2.0.2) databases available at the Center for Genomic Epidemiology website (https://cge.cbs.dtu.dk/services/) (34, 35). For ResFinder, a 90% identity threshold and a minimum length of 60% were used as criteria, whereas for PlasmidFinder, an identity of 95% was utilized.

### Long-read third-generation sequencing

One *K. pneumoniae* isolate per genetic KpnCluster was sequenced using long-read third-generation Nanopore sequencing (18, 36). High molecular weight DNA was isolated using an in-house developed protocol. Bacteria were grown overnight in 1.5 ml Brain heart infusion broth and culture was spun down at 13,000 x *g* for 2 minutes. The pellet was washed and resuspended in 500 µl of 150 mM NaCl. The suspension was spun down at 5,000 x *g* for 5 minutes and the pellet was resuspended in 100 µl of QuickExtract™ DNA Extraction Solution (Lucigen) and 0.1 µl Ready-Lyse™ Lysozyme solution (Epicentre) and incubated for 1 hour at 37°C. Subsequently, 85 µl 10 mM Tris 1 mM EDTA pH = 8 (1x TE), 10 µl proteinase K (>600 mAU/mL, Qiagen) and 5 µl 20% sodium dodecyl sulfate solution were added, and the mixture was incubated at 56°C for 30 minutes. DNA was precipitated overnight at -20°C by adding 0.1x volume 3M sodium acetate pH = 5.2 and 2.5x volume ice cold 100% ethanol. Precipitated DNA was spun down at 13,000 x *g* for 15 minutes and pellets were washed with 1 ml 70% ethanol followed by another centrifugation at 13,000 x *g* for 5 minutes. After drying, the pellet was dissolved in 200 µl 1x TE and diluted to 1 µg with Nuclease-free water.

The Oxford Nanopore protocol SQK-LSK108 (https://community.nanoporetech.com) and the expansion kit for native barcoding EXP-NBD104 was used. Briefly, a shearing step was performed using g-TUBE’s™ (Covaris) to obtain an average DNA fragment size of 8 kb. The DNA was repaired using FFPE and end-repair kits (New England BioLabs) followed by ligation of barcodes with bead clean up using AMPure XP (Beckman Coulter) after each step. Barcoded isolates were pooled and sequencing adapters were added by ligation. The final library was loaded onto a MinION flow cell (MIN-106 R9.4.1). The 48-hour sequence run was started without live base calling enabled on a MinION device connected to a desktop computer. After the sequence run, base calling and de-multiplexing was performed using Albacore 2.3.1 and a single FASTA file per isolate was extracted from the FAST5 files using Poretools 0.5.1 (37). Illumina and Nanopore data were used in a hybrid assembly performed by Unicycler v0.4.4 (38). The resulting contig files were annotated using Prokka and were subsequently loaded into BioNumerics for further analyses (39).

### Minimum Spanning Tree and UPGMA analyses

The BioNumerics software was used to generate a minimum spanning tree (MST) or an UPGMA hierarchical clustering as described previously (16). The MST was based on an in-house *K. pneumoniae* wgMLST scheme. The categorical coefficient was used to calculate the MST. wgMLST clusters were defined as a minimum of two isolates of which the genetic distance between the two isolates was ≤20 genes. An UPGMA clustering of *K. pneumoniae bla*_KPC-2_ and *bla*_KPC-3_ isolates was performed based on the presence and/or absence of antibiotic resistance genes per isolate.

### Plasmid reconstruction by read mapping

The CLC Genomics Workbench version 12.0 software (www.qiagenbioinformatics.com) was used to reconstruct plasmids. For this, complete plasmids obtained by TGS were used as a scaffold to map the trimmed NGS reads of isolates that were from the same genetic wgMLST cluster. A plasmid was scored “present” in an isolate if reads mapped to a reference plasmid of interest and ≥85% of the consensus sequence size in kilo bases was reconstructed. Linear DNA fragments < 5kb were omitted in this study. Nucleotide BLAST analyses on plasmid sequences were performed using the https://blast.ncbi.nlm.nih.gov website and date from October 2019.

### Plasmid content analysis

Bionumerics was used to extract and analyze annotated genes and tranposases in the 18 different plasmids. The data was plotted in Excel. Phaster, the PHAge Search Tool Enhanced Release website (http://phaster.ca/) was used to determine the presence of phage sequences in the plasmids and searches date from October 2019 (40).

### NGS, TGS and plasmid data availability

The Illumina (NGS), Nanopore (TGS) and plasmid sequence data sets generated and analyzed in this study are available in NCBI in the European Nucleotide Archive (ENA) under project number xxx and sequence repositories under Genbank accession numbers xxx. All data supporting the findings of this study are available in this article and its supplementary information files are available upon request.

## Acknowledgements

We thank all the members of the Dutch CPE surveillance study Group and the Dutch medical microbiology laboratories for submitting CPE isolates to the RIVM for the national CPE surveillance program. We thank Dr. Judith W.A. Hoogenboom-Beuving for searching for an epidemiological link of KpnCluster-019 isolates from the Netherlands with the Caribbean. We also thank Prof. Dr. E. Kuijper, Dr. M.G. Mennen and Dr. D.W. Notermans for critical reading of this manuscript.

---

**Supplemental Figure 1. Resistome of *K. pneumoniae bla*_KPC-2_ and *bla*_KPC-3_ non-cluster isolates.** *K. pneumoniae bla*_KPC-2_ isolates were marked blue, and *K. pneumoniae bla*_KPC-3_ were marked magenta. *K. pneumoniae bla*_KPC-2_ and *bla*_KPC-3_ cluster isolates were indicated on the y-axis and AMR genes on the x-axis. The UPGMA clustering was based on the presence (black squares) and absence of AMR genes. Antibiotic classes are indicated above the AMR genes with different colors. A dotted line marks the 85% cut off.

**Supplemental Figure 2. *K. pneumoniae* plasmid gene content (continued).** An UPGMA clustering was performed based on the plasmid DNA sequence for the determination of the genetic relation among the 18 plasmids. Similarity is indicated on the y-axis using a scale from 0 (not similar) to 100% (identical). A similarity of ≥85 to 100% is regarded as the same plasmid. The plasmids are indicated on the x-axis. The presence (black squares) and absence is indicated of annotated genes among the 18 plasmids of six TGS sequenced isolates. If a gene was present twice, blue squares were used and more than two, red squared were used. Colors indicated different clusters of genes with a specific function. The light grey area indicates gene specific content found in only one plasmid.

**Supplemental table 1. Predicted prophage sequences among *K. pneumoniae* chromosomes.**We thank all the members of the Dutch CPE surveillance study Group and the Dutch medical microbiology laboratories for submitting CPE isolates to the RIVM for the national CPE surveillance program. We thank Dr. Judith W.A. Hoogenboom-Beuving for searching for an epidemiological link of KpnCluster-019 isolates from the Netherlands with the Caribbean. We also thank Prof. Dr. E. Kuijper, Dr. M.G. Mennen and Dr. D.W. Notermans for critical reading of this manuscript.

## Members of the Dutch CPE surveillance Study Group

T. Halaby, Analytical Diagnostic Center N.V. Curaçao, Willemstad. R. Steingrover, St. Maarten Laboratory Services, Cay Hill. J.W.T. Cohen Stuart, Noordwest Ziekenhuisgroep, Department of Medical Microbiology, Alkmaar. D.C. Melles, Meander Medical Center, Department of Medical Microbiology, Amersfoort. K. van Dijk, Amsterdam UMC, Department of Medical Microbiology and Infection Control, Amsterdam. I.J.B. Spijkerman, Amsterdam UMC, Academic Medical Center, Department of Medical Microbiology, Amsterdam. D.W. Notermans, Centre for Infectious Disease Control, National Institute for Public Health and the Environment, Bilthoven. J.H. Oudbier, Comicro, Hoorn. M.L. van Ogtrop, Onze Lieve Vrouwe Gasthuis, Department of Medical Microbiology, Amsterdam. A. van Dam, Public Health Service, Public Health Laboratory, Amsterdam. M. den Reijer, Gelre Hospitals, Department of Medical Microbiology and Infection prevention, Apeldoorn. J.A.J.W. Kluytmans, Amphia Hospital, Department of Infection Control, Microvida Laboratory for Microbiology, Breda. M.P.M. van der Linden, IJsselland hospital, Department of Medical Microbiology, Capelle a/d IJssel. E.E. Mattsson, Reinier de Graaf Groep, Department of Medical Microbiology, Delft. M. van der Vusse, Deventer Hospital, Department of Medical Microbiology, Deventer. E. de Jong, Slingeland Hospital, Department of Medical Microbiology, Doetinchem. Anne Maijer-Reuwer, ADRZ medisch centrum, Department of Medical Microbiology, Goes., M. van Trijp, Groene Hart Hospital, Department of Medical Microbiology and Infection Prevention, Gouda. A.J. van Griethuysen, Gelderse Vallei Hospital, Department of Medical Microbiology, Ede. A. Ott, Certe, Department of Medical Microbiology, Groningen. E. Bathoorn, University of Groningen, University Medical Center, Department of Medical Microbiology, Groningen. J.C. Sinnige, Regional Laboratory of Public Health, Haarlem. E. Heikens, St Jansdal Hospital, Department of Medical Microbiology, Harderwijk. E.I.G.B. de Brauwer, Zuyderland Medical Centre, Department of Medical Microbiology and Infection Control, Heerlen. F.S. Stals, Zuyderland Medical Centre, Department of Medical Microbiology and Infection Control, Heerlen. W. Silvis, LabMicTA, Regional Laboratory of Microbiology Twente Achterhoek, Hengelo. J.W. Dorigo-Zetsma, CBSL, Tergooi Hospital, Department of Medical Microbiology, Hilversum. K. Waar, Izore Centre for Infectious Diseases Friesland, Leeuwarden. S.P. van Mens, Maastricht University Medical Centre, Department of Medical Microbiology, Maastricht. N. Roescher, St Antonius Hospital, Department of Medical Microbiology and Immunology, Nieuwegein. A. Voss, Canisius Wilhelmina Hospital, Department of Medical Microbiology and Infectious Diseases, Nijmegen. H. Wertheim, Radboud University Medical Center, Department of Medical Microbiology, Nijmegen. B.C.G.C. Slingerland and H.M.E. Frenay, Regional Laboratory Medical Microbiology (RLM), Dordrecht. T. Schulin, Laurentius Hospital, Department of Medical Microbiology, Roermond. B.M.W. Diederen, Bravis Hospital, Department of Medical Microbiology, Roosendaal / ZorgSaam Hospital Zeeuws-Vlaanderen, Department of Medical Microbiology, Terneuzen. L. Bode, Erasmus University Medical Center, Department of Medical Microbiology, Rotterdam. M. van Rijn, Ikazia Hospital, Department of Medical Microbiology, Rotterdam. S. Dinant, Maasstad Hospital, Department of Medical Microbiology, Rotterdam. M. Damen, Maasstad Hospital, Department of Medical Microbiology, Rotterdam. P. de Man, Sint Franciscus Gasthuis, Department of Medical Microbiology, Rotterdam. M.A. Leversteijn-van Hall, Alrijne Hospital, Department of Medical Microbiology, Leiden. E.P.M. van Elzakker, Haga Hospital, Department of Medical Microbiology, ‘s-Gravenhage. A.E. Muller, HMC Westeinde Hospital, Department of Medical Microbiology, ‘s-Gravenhage. P. Schneeberger, Jeroen Bosch Hospital, Department of Medical Microbiology and Infection Control, ‘s-Hertogenbosch. D.W. van Dam, Zuyderland Medical Centre, Department of Medical Microbiology and Infection Control, Sittard-Geleen. A.G.M. Buiting, Elisabeth-TweeSteden (ETZ) Hospital, Department of Medical Microbiology and Immunology, Tilburg. A.L.M. Vlek, Diakonessenhuis, Department of Medical Microbiology and Immunology, Utrecht. A. Stam, Saltro Diagnostic Centre, Department of Medical Microbiology, Utrecht. A. Troelstra, University Medical Center Utrecht, Department of Medical Microbiology, Utrecht. I.T.M.A. Overdevest, PAMM, Department of Medical Microbiology, Veldhoven. R.W. Bosboom, Rijnstate Hospital, Laboratory for Medical Microbiology and Immunology, Velp. T.A.M. Trienekens, VieCuri Medical Center, Department of Medical Microbiology, Venlo. M.J.H.M. Wolfhagen, Isala Hospital, Laboratory of Medical Microbiology and Infectious Diseases, Zwolle. S. Paltansing, Franciscus Gasthuis & Vlietland, Microbiology and Infection prevention, Schiedam.

